# Multichannel neural spike sorting with spike reduction and positional feature

**DOI:** 10.1101/2022.09.02.506385

**Authors:** Zeinab Mohammadi, Daniel Denman, Achim Klug, Tim C. Lei

## Abstract

Sorting neural voltages measured from a multichannel neural probe to extract the single unit activities of neuronal firing, especially in real-time, remains a significant technical challenge, largely due to the large amount of acquired data and the technical difficulties involved in processing and classifying these neural spikes promptly. Most neural spike sorting algorithms focus on sorting neural spikes post hoc for high sorting accuracy, and reducing the processing time generally is not the chief concern. Here we report on two signal processing modifications to our previously developed single-channel real-time spike sorting (Enhanced Growing Neural Gas) algorithm, which is largely based on graph network. Duplicated neural spikes were eliminated and represented by the neural spike with the strongest signal profile, significantly reducing the amount of neural data to be processed. In addition, the channel from which the representing neural spike was recorded was used as an additional feature to differentiate between neural spikes recorded from different neurons having similar temporal features. With these two modifications, the Graph nEtwork Multichannel (GEMsort) neural spike sorting algorithm can rapidly sort neural spikes without requiring significant computer processing power and system memory storage. The parallel processing architecture of GEMsort is particularly suitable for digital hardware implementation to improve processing speed and recording channel scalability. Multichannel synthetic neural spikes and actual neural recordings with Neuropixels probes were used to evaluate the sorting accuracies of the GEMsort algorithm.

## I. Introduction

Neurons in the brain fire action potentials, also called spikes, which are then processed by multiple hierarchical neuronal circuits to extract biologically relevant information, forming the basis for neural computation. It is therefore not surprising that many experimental approaches include the recording of trains of action potentials in vivo and in some cases even from awake and behaving animals. Such recordings are done by placing a recording channel of a neural probe close to neurons in the extracellular space (extracellular neural recording), and recording trains of action potentials under various experimental conditions(Buzsáki et al. 2017; Rieke et al. 1997; Vogels et al. 2005). One challenge with such recordings is that in many in-vivo extracellular recording conditions, an electrode can pick up neural spikes from several nearby neurons, resulting in so-called “multi-unit” activity (MUA) in the recording trace (Cuevas 2014; Humphrey and Schmidt 1990; Williams 2007). Spike sorting algorithms are then used to separate this multi-unit activity into several sets of “single-unit” activities (SUA), each of which represents the action potential firing pattern of a single neuron (Lewicki 1998; Rey et al. 2015).

Over the years, many neural spike sorting algorithms have been developed, and most of them are focused on sorting neural spikes pre-recorded on a single recording channel (Lewicki 1998; Rey et al. 2015). For instance, several unsupervised classification algorithms, such as K-means, DBScan and WaveClus, have been applied to sort neural spikes based on the classification of the temporal profiles of the pre-recorded neural spikes (Quiroga 2004; Wang et al. 2019). These algorithms mainly focus on improving sorting accuracy and reducing the need for human intervention such as specifying the number of neuronal groups. To that end, we developed an “Enhanced Growing Neural Gas (EGNG)” algorithm for single-channel neural recording based on a graph network approach to learn neural spike clustering features with a few mathematical nodes and edges for classification (Mohammadi et al. 2019a). The advantage of using nodes and edges to form clusters to learn the neural spike feature distribution is to reduce the substantial requirement for computational processing power and system memory storage, while simultaneously achieving rapid neural spike sorting.. This rapid neural spike sorting capability can potentially allow for the sorting of neural spikes in real-time, while neural spikes are being measured from animals during recording sessions.

In recent years, there has been an enormous effort to develop high density neural recording probes, such as Neuropixels, which have hundreds of closely spaced recording channels for simultaneously recording neuronal firing from populations of neurons (Jun et al. 2017; Steinmetz et al. 2021). Due to these technologies, new spike sorting algorithms have been developed is to sort the massive amount of neural spikes recorded from multi-channel neural probes off-line and post-hoc (Chung et al. 2017; Pachitariu et al. 2016; Rossant et al. 2016). Kilosort, for instance, can sort such data based on a template matching and global minimization approach in which the minimization algorithm tries to match the spike templates to the entire neural recording. However, this global minimization approach requires significant computational power and a relatively long computing time to arrive at the sorting outcomes.

Sorting neural spikes and decoding neural information in real-time may open new research paradigms to study neural circuits, such as modulating neuronal spiking activities in a closed-loop feedback manner to alter behavioral responses. To that end, sorting neural spikes in real-time is an important processing step to extract neuronal information on the fly, but this remains a significant technical challenge, especially for handling the large amount of data recorded from high-density multi-channel probes. Compared to sorting pre-recorded neural spikes, real-time spike sorting requires the production of sorting outcomes with very short processing delay (< 10 ms) for closed-loop neuronal feedback applications). Therefore, global minimization approaches cannot result in rapid sorting outcomes. Desirably, the algorithm should only require light-weight computations as well as limited use of memory storage to allow potential hardware miniaturization, which is especially important for implanting the system surgically under the skull. To that end, there have been sustained efforts to develop real-time neural spike sorting algorithms and systems. Many of these real-time spike sorting implementations are mostly based on template matching, where spike templates are calculated by a tethered computer through a short training period and the learned spike templates are then loaded to custom hardware for spike matching (Jun et al. 2017; Park et al. 2017; Wang et al. 2019).

In this paper, we propose a new method – Graph nEtwork Multichannel (GEMsort) - which uses a duplicated spikes data reduction approach to reduce the large amount of neural data, minimizing the required calculation without loss of sorting accuracy. GEMsort also takes advantage of the graph network approach of using nodes and edges to form clusters to dynamically learn the neural spike features. With this graph approach, GEMsort is an inherently lightweight algorithm which can provide rapid neural spike sorting and can potentially be developed with digital hardware for a real-time spike sorting system.

## II. Methods

In this section, the Graph nEtwork Multichannel (GEM) neural spike sorting method for sorting neural spikes measured from a densely spaced multichannel probe is discussed. The GEM algorithm was designed to learn neural spike clustering features dynamically and on the fly and to classify the neural spikes with almost instantaneous sorting outcomes. With this design philosophy in mind, GEMsort can potentially be used to obtain SUAs during neural recording. This information can be used to further decode neural information and may even be extended to modulate neuronal activity in a closed-loop manner. Compared to other spike sorting techniques which minimize the difference between the entire measured data and the reconstructed spike patterns, GEM processes the measured neural spikes only once without the need to retain neural spikes in the system memory, thus significantly saving memory and computational resources. This is largely due to the fact that GEMsort uses movable nodes and interconnecting edges to learn neural spike clustering features and the clusters formed by these nodes and edges are used as a surrogate to represent the clustering relations. Another advantage of using the graph approach is that the clusters formed by these nodes and edges can be adaptively adjusted during the sort to compensate for temporal shape changes during long duration neural recording due to physical movements or other causes.

Figure 1(c) shows the processing pipeline of GEMsort. Neural voltages recorded from a Neuropixels multichannel neural probe were first sent to a noise filter to remove unwanted noise and channel crosstalk contaminations. Neural spikes were then isolated from the neural voltages measured from the multiple recording channels by using a voltage threshold. In contrast to other neural spike sorting algorithms, the neural spike with the highest signal height was identified to represent all the duplicate neural spikes arriving simultaneously to neighboring channels in the closed-spaced recording situation. Features of the representing neural spikes were then extracted using dimensional reduction algorithms and principal component analysis (PCA) in the current implementation. Finally, GEMsort utilizes graph clustering based on the EGNG algorithm in which nodes and edges were used to form clusters to represent the neuron clusters. Another innovation for GEMsort is that the channel from which the representing neuron spikes were measured was also used as a spike feature to allow multichannel sorting, even when the spike features were similar for neurons residing at different locations. The cluster number, spike time and the channel number were estimated as sorting outcomes, which can then be used to construct spike timing histograms for each of the individual neurons.

**Figure 1:**
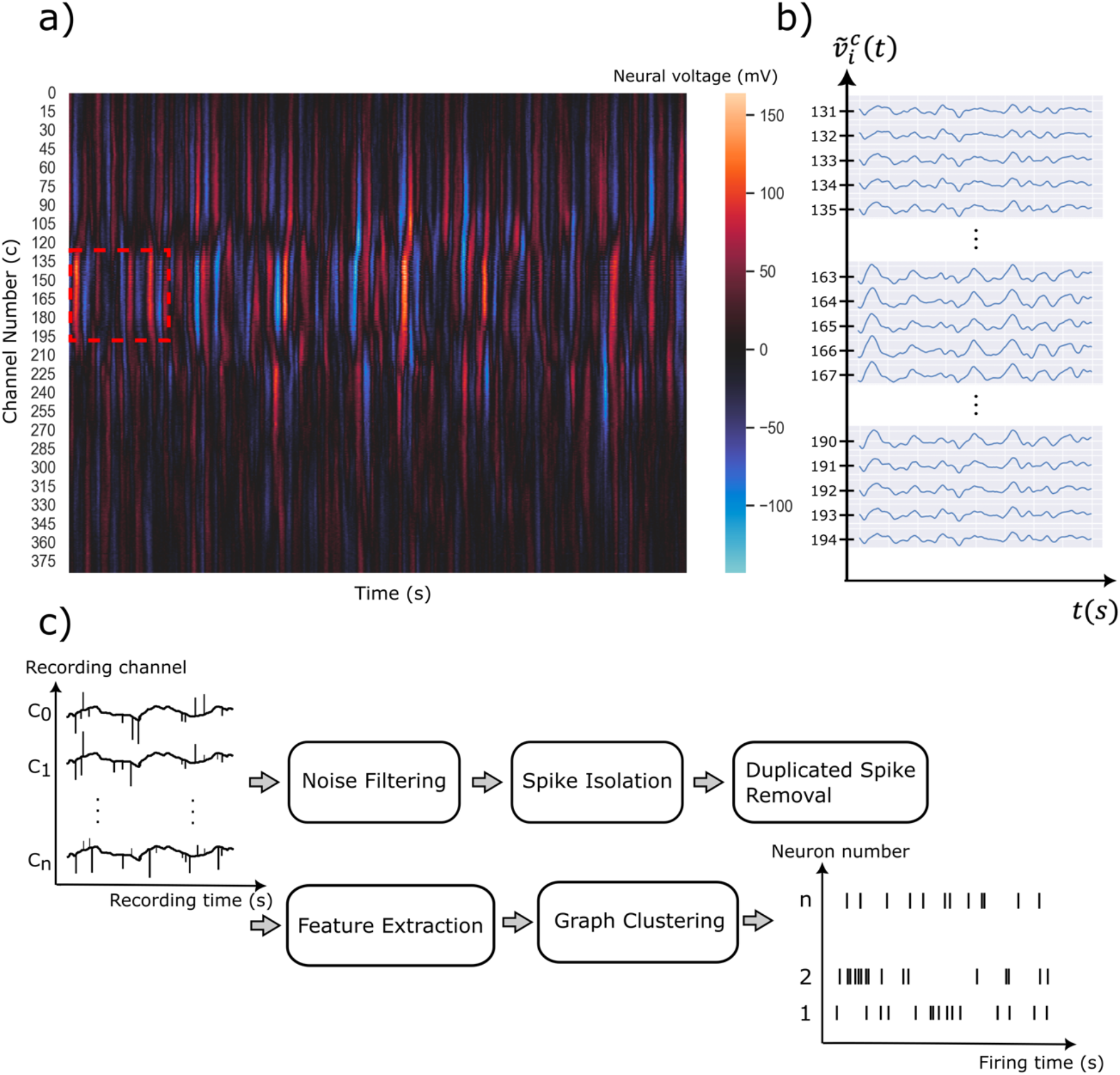
a) Intensity plot of extracellular neural voltage measured by a Neuropixels multi-channel probe recorded near the visual cortical region of a C57BL\6J mouse; b) The expanded voltage plot showing selected channels near the center and the two ends of the red dotted box of a); c) The processing pipeline of the GEMsort spiking sorting algorithm which processes raw multi-channel extracellular voltages to individual neuron spike firing sequence (raster plot).

Figure 2(a) is a schematic diagram illustrating a multichannel neural probe measuring neural firing activities from neurons in the vicinity of the probe. Based on this configuration, the algorithmic details of the GEMsort algorithm will be discussed in the following sections.

**Figure 2:**
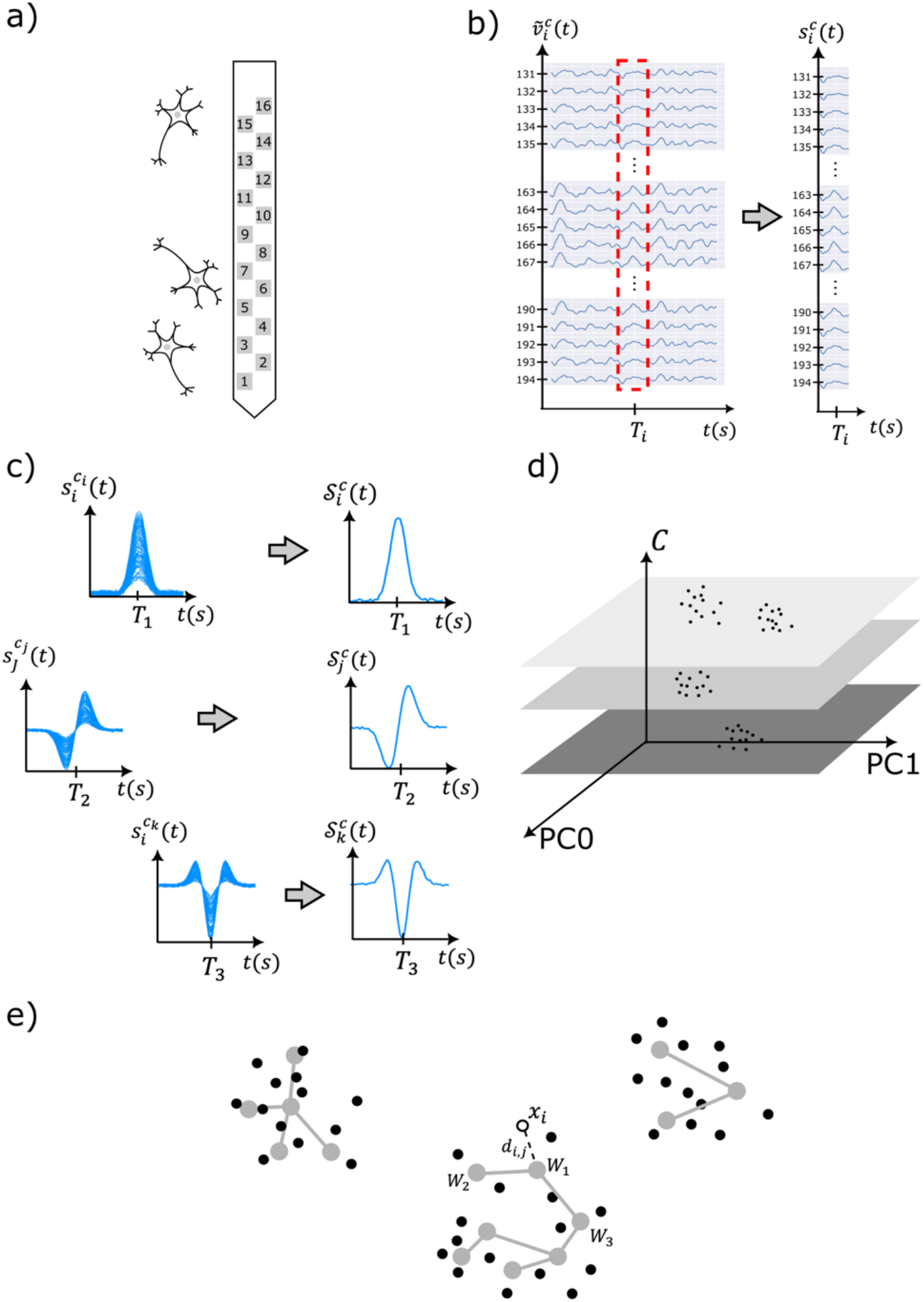
a) An illustration showing a closely spaced multichannel probe for measuring extracellular firings of neurons; b) Extracellular neural voltage 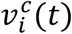 measured from a multichannel probe with channel number *c* being isolated with a threshold value *v*_*th*_ as 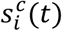; c) multichannel isolated neural voltages 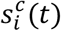 contain duplicated neural spikes and the neural spike with the highest peak voltage was selected as representing neural spikes 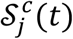; (d) Features of the representing neural spikes 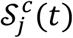 was extracted and projected on the feature space with the channel number *c* used as an additional feature to separate neurons measured from different neural probe channels; (e) Clusters connected with nodes (grey dots) and edges (grey edges) was used in the feature space to learn the neuron cluster distribution. An incoming neural spike can be classified to the closest neuron clusters based on the Euclidian distance *d*_*i, j*_ between the incoming spike and the nodes. The closest node (*W*_1_) and the adjacent nodes (*W*_2_ and *W*_3_) can move closer to the incoming neural spike *x*_*i*_ for adaptable and learning.

### Noise-elimination and neural spike isolation

A second-order high-pass Butterworth filter was employed to remove low-frequency fluctuations and Local Field Potentials (LFP) from the multichannel neural recordings, and a 3dB cut-off frequency of 300 Hz was used to filter the recorded neural voltages. Noise filtered from the neural voltages included electrical and biological artifacts, electrode motion and tissue movements.

A Local Common Average Referencing (L-CAR) filter was then applied to the recorded neural voltages to remove correlated noise common across adjacent channels (Ludwig et al. 2009; Xinyu et al. 2017). The purpose of L-CARS was to remove large-area common noise contaminating the recording channel to allow better spike isolation. For example, assume the neural voltage recorded from the *c*^*th*^ channel of the multielectrode probe at time *t* is *v*_*i*_ (*t*) and *N*_*far*_ and *N*_*near*_ (*N*_*far*_ > *N*_*near*_) are two parameters indicating the numbers of channels to be excluded and included for the noise average. The noise average 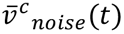 for the *c*^*th*^ channel was then calculated by averaging the neural voltage recorded from channels between *i* – *N*_*far*_/2 and *i*− *N*_*near*_/2, as well as between *i* + *N*_*far*_/2 and *i* + *N*_*near*_/2.

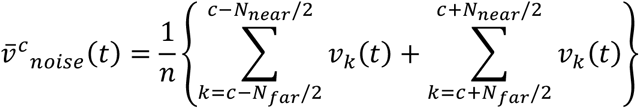

where *n* is the total number of channels included. The L-CARS filtered voltage for channel *c* can be calculated by subtracting this average in which 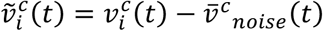.

After noise was removed from the multichannel neural recording, a threshold *v*_*th*_ was used to differentiate and isolate neural spikes from the filtered neural voltage 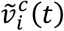, as shown in Fig. 2(b). The threshold *v*_*th*_can be automatically determined based on the standard deviation σ of the multichannel neural recording or manually determined based on custom criteria. In this paper, the threshold was set to be 5 times the signal standard deviation (*v*_*th*_ = 5σ). A neural spike was detected from the filtered neural voltage 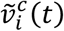 if the signal was higher than the threshold *v*_*th*_ and the isolated neural spike was denoted as 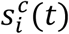 where *i* is the isolated spike index and *c* is the recording channel from which the neural spike was recorded. A spike time *T*_*i*_, which is the peak time of the neural spike, was also associated with the isolated neural spike 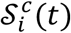.

### Elimination of Duplicate neural spikes

It is apparent from Fig. 1(b) that when a spike was fired by a neuron, it can be simultaneously sensed by multiple neighboring recording channels. This phenomenon is due to the close physical spacing between these channels, which causes the ionic current to be recorded by multiple adjacent channels, effectively leading to the recording of duplicated neural spikes. Generally speaking, the ionic current induced by a neural spike decreases inversely over the propagation distance, causing the recording channel closest to the active neuron to record largest signal strength (Chen et al. 2014; Mohammadi et al. 2019b; Obien et al. 2015; Ranck 1963). In addition, since voltage signal propagation in an ionic solution is an electromagnetic transmission effect which propagates with a speed close to the speed of light, all channels recording the same spike sense the event with no observable signal delays (Humphrey and Schmidt 1990; Jun et al. 2017). This zero-delay property can be used as a criterion to identify duplicated neural spikes.

In this paper, the neural spike with the highest peak voltage was chosen to represent the entire duplicate set of neural spikes, as shown in Fig. 2c). Other selection criteria, such as combining or averaging neural spikes, can also be used to select the representing neural spike; However, our experience indicates that combining these neural spikes did not result in better sorting accuracy. Therefore, in this work we simply use the neural spike with the strongest signal to represent the entire group of duplicated neural spikes for simpler computation. Generally speaking, neural spikes with the highest peak intensities also have the best Signal-to-Noise Ratio (SNR), which typically results in good sorting accuracy (Mohammadi et al. 2019a).

One major advantage of this duplicate spike elimination approach is to significantly reduce the number of neural spikes that need to be processed, hence reducing computation cost. Due to the fast signal propagation speed, only neural spikes with the same peak arrival time were considered to belong to the cluster of duplicate spikes. In addition, channel adjacency is an important factor to consider when determining whether neural spikes arriving the same time belong to the same neural spike fired by a neuron. If the neural spikes had similar temporal features but were measured by channels physically remote, these neural spikes were considered to be fired by two different neurons which happened to have the same temporal features. Pearson’s correlation was then used to determined temporal feature similarity between two neural spikes. The Pearson correlation coefficient *r*_*i,j*_ between the two neural spikes 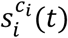 and 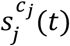 measured from channel *c*_*i*_ and *c*_*j*_ is

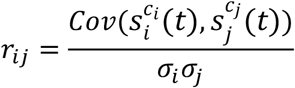

where σ_*i*_ and σ_*j*_, the standard deviation of the *i*^*th*^ and *j*^*th*^ neural neural spikes, and 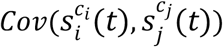 is the covariance between two neural spikes. In this work, the two neural spikes are considered to belong be duplicates if *r*_*i, j*_ is larger than *r*_*TH*_ (*r*_*TH*_ = 0.6 was used in the work). Once neural spikes with high correlations were identified, the neural spike with the highest peak signal was selected as the representing neural spike 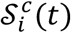 (*c* as the channel measuring the strongest neural spike). The remaining duplicated neural spikes were discarded since including these spikes in the classification process can lead to counting duplication.

### PCA based feature extraction trained with initial spikes

Once the representing neural spike 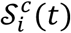 was determined, the neural spike was then sent to a dimension reduction algorithm for feature extraction. In addition, the representing neural spike 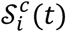 was associated with a channel feature *c*, which was used to differentiate neural spikes measured from different areas of the neural probe.

PCA was used to extract features from the representing neural spike 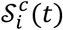. Traditional PCA analyzes the entire set of neural spikes to determine the principal components for feature extraction, but this approach is incompatible with real-time neural spike sorting. Since neural probes are mounted securely within the animal brain, temporal profiles of the recorded neural spikes do not change rapidly, but can slowly drift after hours of recording due to small movements of tissue. Based on this assumption, principal components *f*_*n*_ (*t*) can be learned merely by using neural spikes measured during a short training period based on the standard PCA routine. Features of streamed-in neural spikes can be extracted using the estimated principal components. The *n*-th features 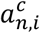 of a representing neural spike 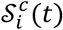 measured from channel *c* can be calculated by

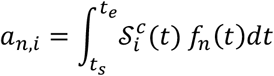

where *f*_*n*_ (*t*) is the *n*-th principal component; *t*_*s*_ and *t*_*e*_ are the beginning and end time of the isolated neural spike epoch. In GEMsort, the channel *c* is used as one of the features to separate neural spikes measured from different locations. The feature *c*_*i*_ of the representing neural spike 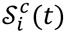 is *x*_*i*_ = {*c*_*i*_, *a*_0,*i*_, *a*_1,*i*_, *a*_2,*i*_⋯}. This channel feature *c* is an important improvement to the original single channel algorithm to allow for the separation of clusters measured from different recording channels to avoid cluster overlaps, as shown in Fig. 2(d). Due to dimensional reduction of PCA, only the first few features of neural spikes (*n* = 2 for this work) are needed to be sent to the next step for clustering.

### Graph Network based neuron spike clustering

Neural spikes fired by the same neuron have similar temporal features, which allows cluster formation in the feature space. Unsupervised learning techniques can then be used to classify these clusters. In GEMsort, instead of examining the entire set of recorded neural spikes, a graph network approach allows for the examination of neural spikes one at a time, and this individual spike examination allows for rapid sorting. In addition, the algorithm learns the neural clustering distribution on the fly through forming, deleting or moving these surrogated clusters constructed by interconnecting nodes with edges.

We have previously demonstrated real-time graph-based spike classification with neural spikes recorded from a single recording channel (known as Enhanced Growing Neural Gas classification) and the details of this single-channel classification algorithm can be found in Ref. (Mohammadi et al. 2019a). Here, a brief description of that method is provided. In EGNG, neural clustering can be learned by interconnected nodes with edges forming mathematic clusters in the feature space. When a neural spike is measured, the Euclidean distance between the spike features and the nodes are compared and the neural spike is classified to the cluster group containing the node closest to the neural spike. Suppose the features of the representing neural spike 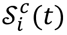 is *x*_*i*_ = {*c*_*i*_, *a*_0,*i*_, *a*_1,*i*_, *a*_2,*i*_⋯} and the *j*^*th*^ mathematical node is denoted as *N*_*j*_, the Euclidean distance between the representing neural spike feature 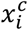 and the *j*^*th*^ mathematical node *w*_*j*_ is

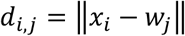

Edges connecting the nodes to form clusters can be created by identifying the closest node 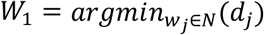 and the second closest node 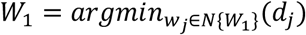 to the neural spike. If the two closest nodes have not been connected by an edge, a new edge is formed to connect the two nodes. Through these edge connections, clusters can be formed for classification. In addition, if the two nodes connecting an edge have not been identified as closest nodes for an extended period, the edge can be deleted to indicate the dissociation of the two nodes, eventually leading to cluster dissociation.

This node and edge approach allows the clusters to adapt and learn the neural cluster distribution by moving the closest node *W*_1_ and its neighboring nodes *W*_*j*_ (*j* ≠ 1) closer to the neural spike features based on the moving rates (*e*_*S*1_ and *e*_*nbr*_).

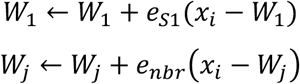

New nodes can be procedurally added to expand the clusters when neural spike features repeatedly appear in an area, and unused nodes can be deleted when neural spikes are no longer being recorded in certain regions in the feature space. Based on this dynamic clustering, clusters can dynamically adjust, formed and deleted when the neural clustering distribution is slowly changing during long recording sessions.

### Rapid neural spike clustering through identifying the closest cluster

With this graph approach, the neural clustering distribution can be adaptively learned by the interconnecting nodes and edges, retaining previously recorded neural spikes in the system memory to inform that future clustering is not needed. Classification of a neural spike can be rapidly estimated through determining the graph cluster containing the closest node *W*_1_ to to the neural spike. The output of GEM spike sorting contains the neuron cluster number, the peak neural spike firing time and the channel feature. This information can be used for further neural data analysis. For instance, a firing time histogram against the neuron cluster number can be constructed based on this sorting outcome.

The employed parameters for the Multielectrode EGNG algorithm are shown in Table 1.

**Table 1.** Parameters required for the GEMSort spike sorting algorithm. Typical parametric range and the parameter values used for the results demonstrated in this paper are listed in the first and second columns respectively.

| Parameters | Typical ranges | Values used in this paper |
| --- | --- | --- |
| Maximum node count ($N_{\text{EGNG}}$) | 3 to Max (it depends on the data and the recording area of the brain) | 10 to 15 for every channel of the synthetic 16-channel MEA (it can be changed based on the Neural density of the recording area) |
| Edge pruning threshold ($a_{\text{th}}$) | A fraction of total node count (e.g., half or one-third). | 4 to 8 for the synthetic 16-channel MEA data |
| Number of initial nodes ($N$) | 2 | 2 |
| Moving rate of $W_1$ ($e_{s_1}$) | (0, 1) | 0.8 |
| Moving rates of $W_j$ ($e_{\text{nbr}}$) | (0, 1), $e_{s_1} > e_{\text{nbr}}$ | 0.001 |
| Insert parameter reduce rate of $w_e$ and $w_e'$ nodes ($\alpha$) | (0, 1) | 0.5 |
| Insert parameter reduce rate of other nodes ($\beta$) | (0, 1) | 0.01 |
| Number of iterations before inserting a new node ($\lambda$) | 5 to 50 | 10 |

## III. Results

The GEMsort algorithm was implemented using Python version 3.7 programming language with Jupyter notebook and was run on a 3.4 GHz i7-4770 desktop computer with no use of GPU computation. Two sets of multichannel neural extracellular recording data were used to evaluate the performance of the GEMsort algorithm. The first set of data was a 16-channel synthetic neural data, and the second set of data was a pre-recorded electrophysiology neural recording measured with a Neuropixels probe.

### Performance evaluation using a synthetic 16-channel neural dataset

#### Generation of the synthetic spike data

Multichannel synthetic neural data was useful to evaluate the performance of the GEMsort algorithm due to the flexibility of adjusting the parameters of the dataset to achieve certain testing conditions while maintaining reasonable levels of complexity and realism. To imitate spontaneous neuronal background firing which generally follows Poison statistics, the extracellular spike sequences of the 16 channels were generated with a Poisson generator. User selected firing rates *f*_*rate*_ (1 < *f*_*rate*_ < 200) were used to constrain the Poisson generator to generate firing sequences for the neurons for specific firing rates. Eight firing neurons were then placed in the immediate vicinity of channel 0, 3, 5, 7, 9, 11, 13 and 15, causing these channels to receive the strongest neural signals from their respective closest neurons. Neural spikes were generated using different wavelet shapes to mimic various temporal profiles fired by different neurons. The temporal profiles of the neural spikes used for the 8 neurons included Gaussian, Richer, Biorthogonal, and Daubechies wavelets (Lee et al. 2019). Gaussian noises with a variance of 0.02 were added to the synthetic neural spike data to simulate noise contamination caused by the recording environment or other sources. Signal reduction due to distance (*r*) between the neurons and the recording channels were modeled based on a parabolic voltage potential *V*_*e*_(*r*) = 1/4πσ*r*, where the extracellular fluid conduction σ = 0.35 [36]. Figure 3(a) shows an example of syntenic neural voltages plotting the 16 channels to mimic a multi-channel neural recording.

**Figure 3:**
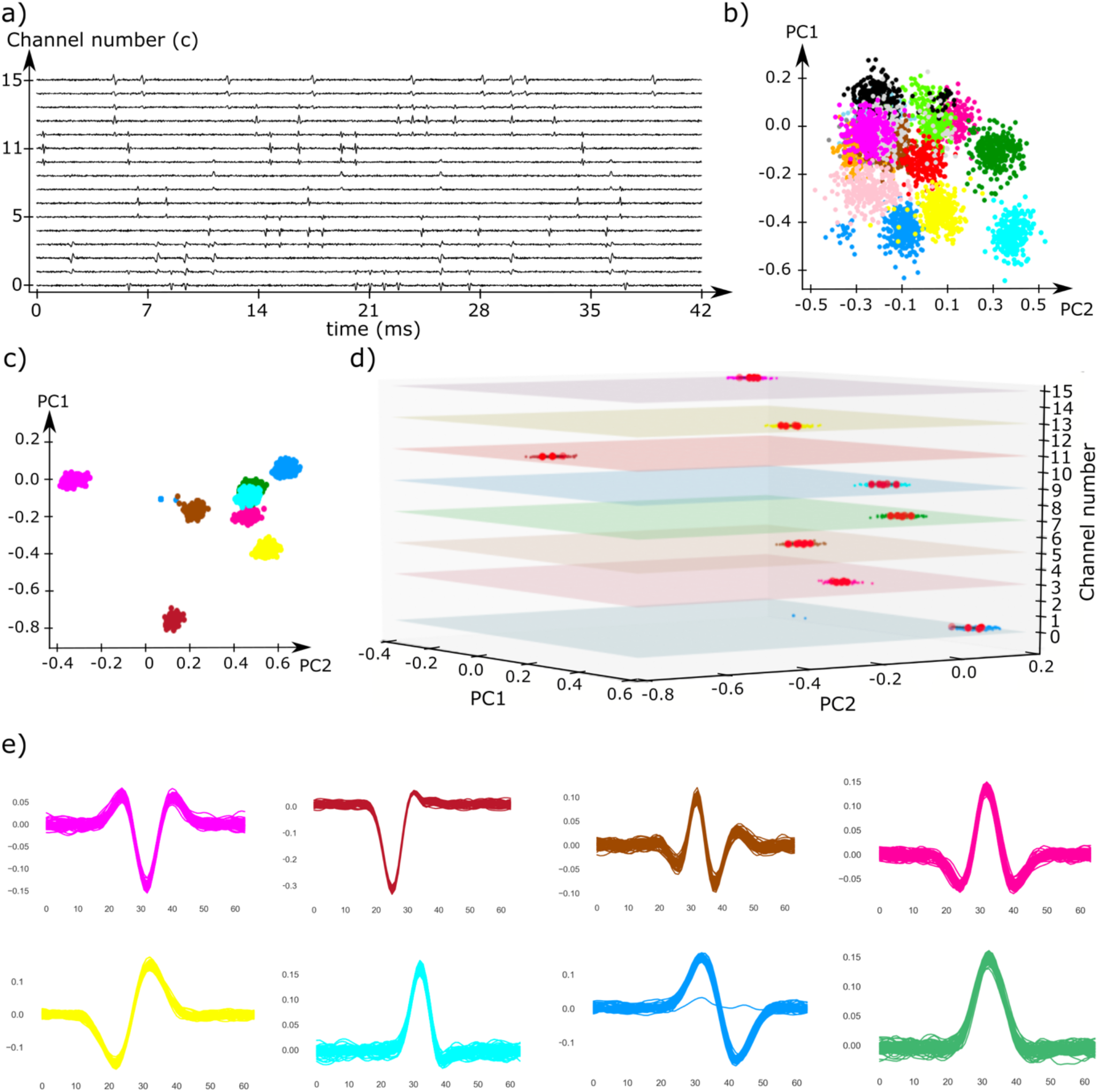
a) Simulated neural voltages of 16 neural recording channels mimicking neuronal firing of 8 neurons; b) Feature representation of all neural spikes measured by the 16 recording channels without elimination of duplicated neural spikes. Significant cluster overlap can be observed due to the artificial noise added to the neural spikes; c) Feature representation of only the representing neural spikes after duplicated neural spike elimination. Neuronal clusters were more separated from one another despite there are still some clustering overlaps (red, cyan and green) due to similar spike profiles; d) The overlapping clusters can be separated by adding the channel position number as an additional feature to separate neural spikes with similar temporal profiles but the firing neurons were distributed at different channels; e) The single unit activities (raster plot) of 8 firing neurons can be reconstructed based on the sorting outcomes of GEMsort.

#### Sorting synthetic neural spikes with GEMsort

The 16-channel synthetic neural voltages to mimic neural spike firing by 8 neurons were sent to the GEMsort algorithm through the processing pipeline for neural spike sorting. As discussed in the methods section, for closely spaced multichannel neural probes, the same neural spikes can be picked up by several adjacent recording channels, resulting in the same spike being sorted multiple times. Figure 3b) shows the feature representation of all neural spikes recorded from the 16 recording channels without any duplicated neural spike elimination. It is obvious that the neural spikes were overlapping with one another. The spreading and overlap of the clusters was due to artificial noise being added to the neural spikes and these overlaps make spike classification difficult. In addition, if all the neural spikes were sorted without elimination, the same neural spikes were sorted multiple times, leading to incorrect spike rates for sorting outcomes. In contrast, Fig. 3c) shows the feature representation of only the representing neural spikes. With the elimination of the duplicated neural spikes and only using the strongest neural spikes for the sort, the clusters were more separated from one another, making cluster classification much easier and more accurate. Another important benefit of only sorting the representing neural spikes is that it avoids over-sorting the duplicated spikes such that accurate firing rates can be calculated for further signal analysis. However, there are still clusters (red, cyan and green) which closely overlap in Fig. 3c) since the neural spikes of these clusters had similar temporal profiles, and therefore similar features. GEMsort solved this issue using the recording channel number as an additional feature to separate neural spikes with similar temporal profiles but fired from different neurons at different locations on the neural probe. Figure 3d) is the 3D feature plot using the channel number as a feature to separate overlapping clusters. With the additional channel feature, the overlapping clusters were further separated to allow proper neural spike classification. Over the total 2242 isolated neural spikes, without the channel feature, the sorting accuracy was only 79.8% due to the significant cluster overlap. With the additional channel feature, the sorting accuracy was improved to 93.0%, and the sorted neural spikes of the 8 cluster groups were plotted in Figure 3e). The remaining misclassification was likely due to closely spaced neurons firing at similar time intervals, resulting in temporal profiles that cannot fall into any of the cluster groups leading to misclassification. Since the spike time *T*_*i*_ was associated with the representing neural spikes 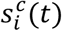,once the neural spikes were sorted, the sorting outcomes could use *T*_*i*_ to construct a raster plot. Figure 3f) is an example raster plot generated with the sorting outcomes of this specific simulated neural voltages.

Figure 4 illustrates the classification process using a graph network. In the beginning of the sort, as shown in Fig. 4a), only a small number of neural spikes were streamed into the feature space, and a few mathematical nodes were generated to cover these initial spikes to form clusters. As more neural spikes were streamed into the feature space and analyzed by the algorithm, more nodes were generated and interconnected to form larger clusters as shown in Fig. 4b, allowing the sort to be more accurate as the cluster stabilized. At the end of the sort, as shown in Fig. 4c), the clusters were fully stabilized to cover the neural spikes for accurate sort. Note that these clusters could continue to adapt, should the temporal shapes of the neural spikes change, allowing the algorithms to learn the neural spike distribution on the fly to accustom for changes or noise contaminations. This self-learning feature makes GEMsort particularly useful for sorting neural spikes for long-duration neural recording. It is also important to mention that as the streamed neural spikes were analyzed and learned by the graph clusters, the neural spikes were not further retained in the system memory, allowing GEMsort to be particularly nimble and not requiring extensive system hardware. A movie demonstrating the classification process of the graph network classification approach can be found in the supplementary information, illustrating how graph networks can learn and adapt to neural features and graph clustering on the fly while neural spikes were streamed into the processing pipeline.

**Figure 4:**
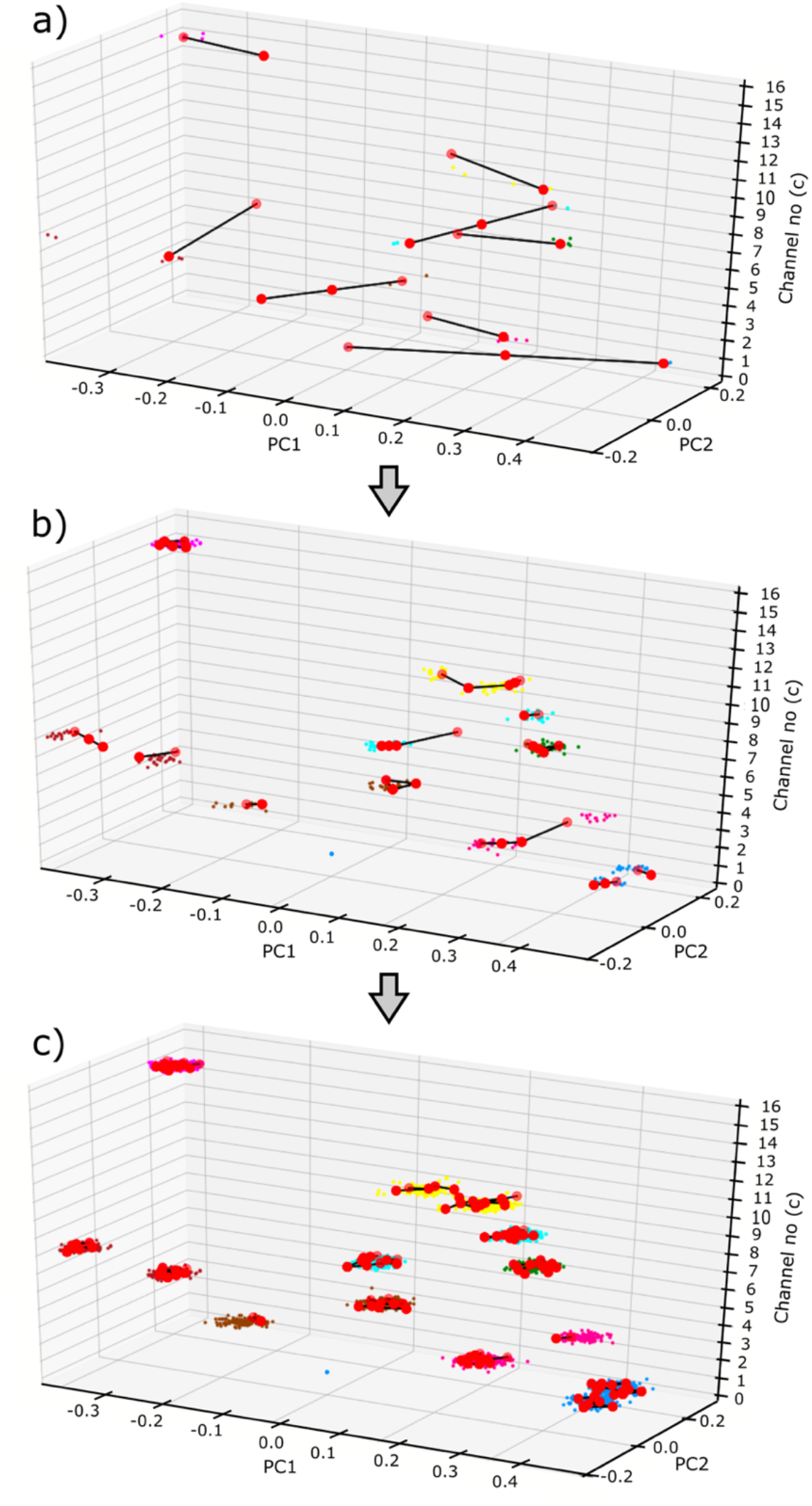
Neural spike classification in the feature space. a) Early classification process when only a few neural spikes streamed into the algorithm and early interconnecting nodes were formed to learn the neuronal clustering distribution; b) As more neural spikes were streamed into the algorithm, more nodes were generated to form larger clusters and the neural spikes started to be correctly sorted; c) Through node movements and edge connections or eliminations, stable neuron clusters can be formed to provide rapid neural spikes sorting. If feature distribution of the neuron clusters changed, the nodes and edges can move to conform to the new feature distribution to allow continue proper neural spike sorting.

### Sorting multichannel neural voltages measured by Neuropixels

#### Neural data recorded by a Neuropixels probe

The GEMsort algorithm was also evaluated using prerecorded multichannel neural data measured by a Neuropixels probe. Neuropixels is a single shaft multi-channel neural probe consisting of 384 recording channels selectively measuring from 960 recording sites with a spacing between the recording sites of ∼20 μm. For the neural data used in this paper to test the GEMsort algorithm, neural data were recorded by Neuropixels Phase 3A prototype probes (Jun et al. 2017) in awake mice, following procedures described in detail elsewhere (Denman and Reid 2019). All animal procedures were approved by the Allen Institute for Brain Science Institutional Animal Care and Use Committee (IACUC). Mice in this study were male C57BL\6J aged 60 – 182 days. Briefly, animals were first outfitted with a permanently attached head fixation device and habituated to the recording apparatus. On the day of recording, an approximately 2×2 mm cranial window was opened over the primary visual cortex, and a Neuropixels probe lowered through cortex, subiculum or hippocampus, and ventral thalamic structures. Probes were inserted with piezoelectric manipulators at 50 - 100 μm /minute to their target depth. Increased neural activity was elicited with visual stimuli, which consisted of full-field flicker, repeated presentation of natural images, and a repeated naturalistic movie clip.

#### Sorting comparison between GEMsort and Kilosort

A 200 second segment of Neuropixels neural recording data was sent to our Python GEMsort code and the Kilosort spike sorting routine to compare sorting results. The sampling rate of the recording was 30 kHz/s which resulted in 6,000,000 data points for each recording channel. With the 384 total recording channels, a total of 384×6,000,000 data points were processed by both algorithms. A Windows 10 desktop computer equipped with a i7-4770 CPU running at 3.40GHz, a Nvidia Titan RTX GPU, and 24 GB of system memory were used for both algorithms. Despite not using any GPU processing for GEMsort, the processing time for sorting 200 seconds of 384 recording channels was ∼279 s. In comparison, the processing time for Kilosort was ∼543 s, which was approximately two times longer than GEMsort. In addition, Kilosort requires manual curation to combine or split clusters, which is not required for GEMsort, and this manual processing time was subjective and therefore was not included.

Figure 5 shows the confusion matrix for comparing the sorting outcomes between GEMsort and Kilosort. The left-vertical and the top-horizontal axes show the GEMsort cluster ID number and the Kilosort cluster ID number respectively. The square element within the confusion matrix represents the numbers of common spikes between the two sorting algorithms. A grey-scale intensity within the square element was used to indicate the matching percentage between the two sorting algorithms (the darker of element, the better the match). The elements of the bottom-horizontal axis show the numbers of neural spikes and the percentages (in grey-scale intensities) that could be sorted to the corresponding GEMsort clusters but could not be matched to any Kilosort clusters. Similarly, the right-vertical axis shows the opposite situation. For this Neuropixels recording clip, Kilosort detected 512 neuronal clusters before manual curation. After manual curation, 93 selected clusters remained based on visual inspection and labeling the clusters as valid, multi-unit activity, or noise contaminations, as well as merging or splitting the clusters. By comparison, GEMsort detected 106 neural clusters without the need for manual curation. Based on the sorting results, the GEMSort and Kilosort neuron IDs were respectively numbered from cluster 1 to 93 and from cluster 1 to 106.

**Figure 5:**
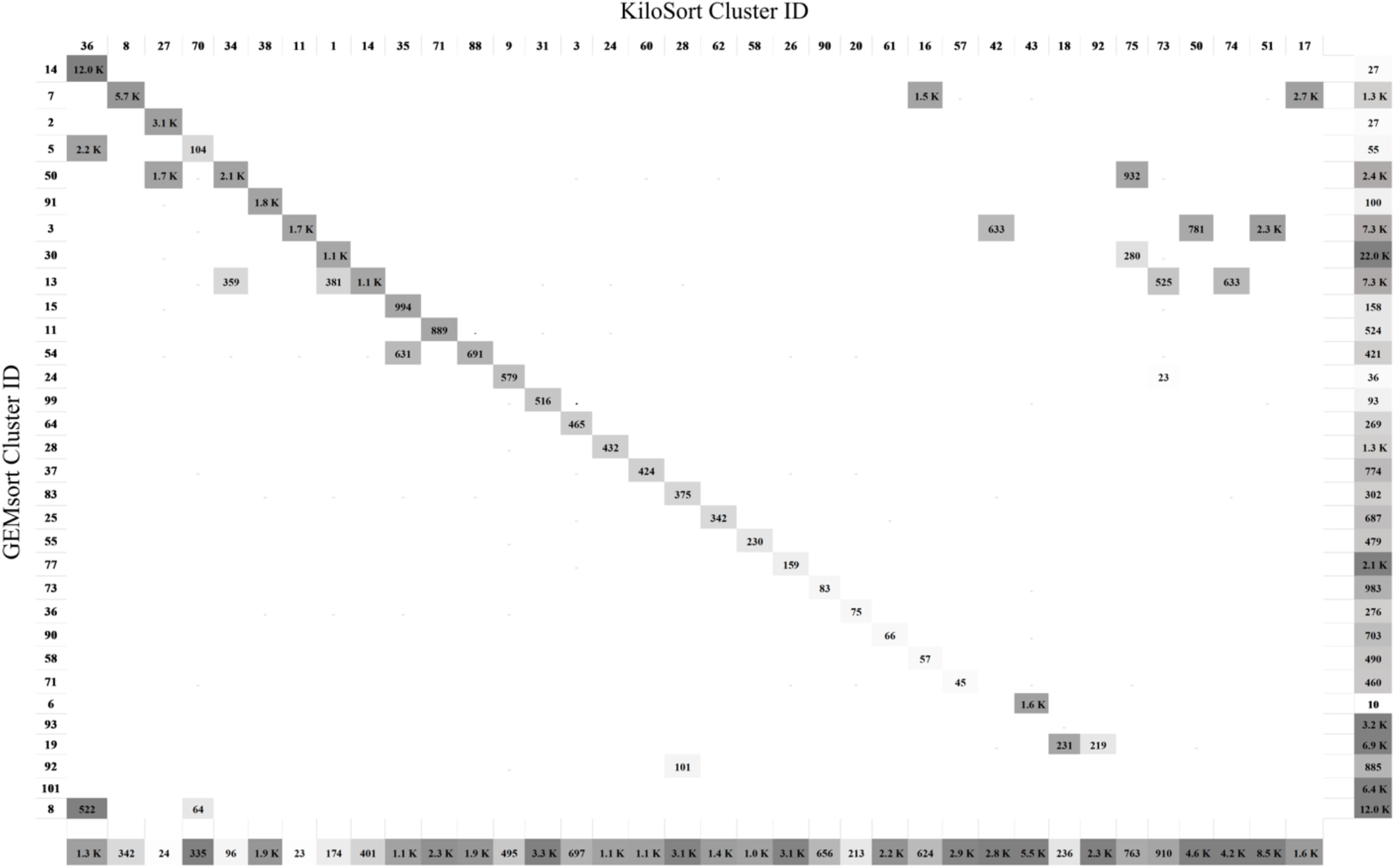
Confusion matrix comparison of sorting agreement between GEMsort and Kilosort. The left-vertical and the top-horizontal axes show the GEMsort cluster ID number and the Kilosort cluster ID number respectively. The square element within the confusion matrix represents the numbers of common spikes between the two sorting algorithms and a grey-scale intensity within the square element were used to indicate the matching percentage between the two sorting algorithms. The elements of the bottom-horizontal axis show the numbers of neural spikes and the percentages in grey-scale intensities that could be sorted to the corresponding GEMsort clusters but could not be matched to any Kilosort clusters.

The diagonal elements of Fig. 5 indicate matched sorting outcomes between the two algorithms. Besides these matched results, there are a few off-diagonal elements which indicate sorting discrepancies between the two algorithms. These discrepancies were mainly caused by one algorithm which considered the neural spikes to belong to the same neural cluster while the other algorithm considered them to belong to two different clusters. The neural spikes on the bottom-horizontal and right-vertical axeswere considered to be neural spikes by one algorithm but considered as noise or MUA by the other algorithms. This discrepancy may be caused by the subjective judgement in the manual curation process or different noise detection thresholds used for both algorithms.

## IV. Discussion

GEMsort was designed to sort neural spikes rapidly. Compared to many existing neural spike sorting algorithms which were designed to sort neural spikes post-hoc in an off-line manner, GEMsort uses a different approach to analyze the neural spike only once and estimate the neuronal clustering with a short processing time. In this paper, the algorithm was implemented using Python as a software routine and therefore requires some processing time to sort the neural spikes. We are currently working on a digital hardware implementation of the GEMsort algorithm to allow sorting neural spikes with real-time processing speed with millisecond-level processing delay. In addition, GEMsort allows several of its processing steps, including the spike elimination and classification modules, to be implemented using a parallel computation architecture. This modular capability allows scaling up the number of neural spikes to be processed without scarifying the processing time. This parallel design is particularly suited to be implemented by digital hardware but may be difficult to implement with a software approach due to the limited number of processing cores allowed in CPUs and the inflexibility of GPU computations.

All the parameters needed for the GEMsort algorithm are listed in Table 1. More nodes were used for the Neuropixel data, but otherwise the same parameters were used to sort both the 16-channel synthetic neural data and the Neuropixels data. In addition, GEMsort does not require any manual intervention in the sorting. Not only can this avoid subjective sorting judgements, this feature can also be important for close-loop neural circuit control or live and real-time neural data analysis during an experiment, since there might not be time to perform manual processing, especially when an animal is under sedation and only limited amount of time is available to perform an experiment (Edward et al. 2018).

In this paper, the sorting results between GEMsort and Kilosort were compared. This comparison was used mainly to evaluate whether GEMsort can reach the sorting performance of Kilosort in a simulated off-line sorting scenario. Despite a high degree of agreement between the two sorting algorithms, there remains some discrepancy, largely caused by disagreement as to whether the neural spikes belong to the same cluster group or not. In addition, there were also disagreements regarding whether some neural spikes should be considered as noise or MUA. In Kilosort, a manual curation step is required to aid this determination process. For GEMsort, further refinement of the algorithm and the possibility of adding human pre-determined criteria can help guide the algorithm to do better clustering determination.

GEMsort currently cannot correctly sort mixed neural spikes in situations where two nearby neurons fired at the same time, and considered these overlapping events either as outliers or misclassified them to a neuronal cluster. In Kilosort, a second processing pass was used to identify these overlapping neural spikes by matching them to mixed spike templates estimated from the initial processing pass. GEMsort can utilize the same idea by adding a final processing stage at the very end to match neural spikes that are not close to the neuron clusters to two mixed cluster groups. However, this idea has not been implemented in the current version of the algorithm and will be explored in the future. On the other hand, it is well known that neuronal firing and neural signal encoding in the brain has significant firing variability. Neuronal responses to the same stimulus typically vary to some degree on a trial by trial basis. Therefore, firing rate averages acquired from many trials are typically used to study neuronal circuits. However, the brain uses coding redundancy to combat these fluctuations and uses ensemble coding or other more complex encoding schemes. Therefore, for real-time neural information decoding, small spike sorting fluctuations due to occasional spike overlaps may not be a significant factor in neuronal decoding interpretation. The lack of real-time neural signal decoding tools is one of the reasons why single trial real-time neural signal has yet to be used for neuroscience studies, and future neural signal processing should be further developed to solve the problem. To that end, GEMsort may be a viable option for extracting real-time SUA to allow for the exploration of neural circuits from a new angle.

## V. Conclusion

Through the combination of duplicated spike elimination, positional feature and graph network clustering, GEMsort demonstrated that it has the potential to be used to sort neural spikes measured in real-time. The hardware requirement for the GEMsort algorithm is also significantly less than some of the existing neural spike sorting routines. These technical benefits make GEMsort aviable choice as the frontend neural spike sorting processing unit to extract SUAs for real-time neural information decoding.

## Notes

### Competing Interest Statement

The authors have declared no competing interest.

